# Advantages of outcrossing in *Plasmodium falciparum*: insights from genetic crosses using fluorescent labelled parasites

**DOI:** 10.1101/2025.07.18.665596

**Authors:** Xue Li, Kathrin S. Jutzeler, Biley A. Abatiyow, Nastaran Rezakhani, Meseret T. Haile, Amanda S. Leeb, Hardik Patel, Stefan H.I. Kappe, François Nosten, Ian H. Cheeseman, Michael T. Ferdig, Timothy J. C. Anderson, Ashley M. Vaughan, Sudhir Kumar

## Abstract

Malaria parasites are obligately sexual hermaphrodite protozoans with gamete fusion occurring in the mosquito midgut, followed by meiosis and recombination. Malaria parasite populations show a spectrum of populations structures ranging from predominantly selfing to highly outcrossed. However, the fitness consequences of selfing and outcrossing for malaria parasites are poorly understood. This project was designed to investigate the dynamics of gamete fusion within the mosquito midgut and the relative fitness of selfed and outcrossed zygotes. We generated florescent-labelled clones of NF54 (mCherry), an African parasite, and NHP4026 (GFP), a Thai parasite, crossed these parasites, and scored genotypes of 8540 oocysts from 435 mosquitoes sampled from 7 to 14 days post infection. We observed decreasing proportions of outcrossed oocysts and increasing levels of inbreeding over the course of the infection in two independently replicated crosses. These results are consistent with the faster maturation of transmissible sporozoites derived from outcrossed compared with selfed oocysts. Our results suggest a substantial outcrossing advantage, perhaps because this allows for the removal of deleterious mutations accumulated during asexual parasite replication in the vertebrate host. We also found that selfed NF54 oocysts were larger than outcrossed or selfed NHP4026 oocysts, which may influence production of sporozoites and onward transmission. We conclude that fluorescent labelled parasites provide clear resolution of mating patterns, temporal dynamics and transmission potential of malaria parasites in mosquitoes. Importantly, faster maturation of outcrossed parasites can maximize levels of recombination in transmitted malaria parasite populations.

## INTRODUCTION

Many animals and plants are simultaneous hermaphrodites that produce both male and female gametes and have the capacity for both inbreeding and outbreeding (Goodwillie, et al. 2005; Jarne and Auld 2006). Self-fertilization has obvious benefits, because reproduction is possible when partners are not available (reproductive assurance). Furthermore, the offspring are 100% related to parents. However, inbreeding can also result in accumulation of deleterious mutations and declining fitness (Charlesworth and Willis 2009). Outcrossing allows purging of deleterious mutations and restoration of fitness, but offspring share only 50% relatedness to their parents. Nevertheless, the advantages of outcrossing are evident from the elaborate mechanisms used by many hermaphrodite plants to avoid inbreeding, including self-incompatibility and physical barriers to pollen transfer (Charlesworth 2006; Zhang, et al. 2024; Shang, et al. 2025). Furthermore, many controlled experiments have directly compared fitness of inbred and outbred plants and animals, revealing substantial fitness advantages to outbreeding (Willis 1993; Coltman, et al. 1999; Meagher, et al. 2000; Christen and Milinski 2003; Goodwillie, et al. 2005).

Malaria parasites are obligately sexual hermaphrodite protozoans that produce both male and female gametes. These parasites have a mixed mating system, and the contribution of inbreeding and outbreeding varies dramatically in different locations (Anderson, Haubold, et al. 2000). Haploid blood stages differentiate into immature male or female sexual stages (gametocytes) in the blood stream. These are ingested by mosquitoes and male and female gametes mature, fuse in the mosquito midgut to form a diploid zygote which develops into a short-lived motile stage, the ookinete, where meiosis and recombination occur. The ookinete crosses the mosquito midgut and attaches to the basal lamina and dedifferentiates into an extracellular oocyst which contains the four haploid products of meiosis. These divide mitotically and after 10-16 days the oocyst ruptures, releasing one to five thousand (Rosenberg and Rungsiwongse 1991) haploid sporozoites into the hemolymph which ultimately invade the salivary gland. Salivary gland sporozoites can be transmitted to the next vertebrate host when the mosquito takes a bloodmeal. During fertilization, if male and female gametes are from the same parasite genotype – which can happen when a mosquito feeds on a person infected with a single parasite clone – then gamete fusion is inbred. While meiosis and recombination occur, there is no change in parasite genotype from the parent. On the other hand, when gametocytes of two or more genotypes are present in the blood meal, both inbreeding and outcrossing can occur. In the case of outcrossing, recombination results in the reshuffling of chromosomal segments from different parental parasites and the formation of recombinant progeny.

Several controlled genetic crosses have been conducted in the human malaria parasite *Plasmodium falciparum*. This is achieved by growing parasite clones in *in vitro* sexual culture conditions and allowing mosquitoes to feed on blood containing male and female gametocyte stage parasites from two different clones. Vertebrate hosts are needed for mammalian stage parasite development and splenectomized chimpanzees were initially used (Su, et al. 2007). Genetic crosses are now conducted using humanized mice seeded with human hepatocytes that support parasite liver stage development and transition to asexual blood stage replication (Vaughan, et al. 2015; Vendrely, et al. 2020; Button-Simons, et al. 2021; Mok, et al. 2023). Cloning and sequencing of haploid parasites from the mammalian host from prior crosses conducted in chimpanzees or humanized mice have frequently revealed a surprising deficit of inbred parasites (Ranford-Cartwright, et al. 1993; Button-Simons, et al. 2021; Amambua-Ngwa, et al. 2023). For example, genotyping of 376 blood stage cloned progeny derived from three independent *P. falciparum* crosses between NF54 and NHP4026 (or NF54^mCherry^ and NHP4026^GFP^) recovered only six selfed progeny (1.6%) (**Table 1**). Assuming random mating and equal frequencies of gametes from the two parents, the expectation would be to recover 50% of inbred parasites (25% of each parental genotype); when gamete frequencies are unbalanced frequencies of inbred parasites should be >50%. Preferential outcrossing of gametes in the mosquito midgut, or selection of outcrossed parasites during development may explain the deficits of selfed parasites observed.

**Table 1.** Summary of previous genotyping of cloned blood stage progeny. Cloned progeny were sequenced as part of the Malaria Crosses Collaboration project (https://mcc.nd.edu/) and sequences are available in GitHub repository (https://github.com/emilyli0325/P01-cloned.progeny) (Li, et al. 2022). Progeny were classified as follows: (1) those with ≥90% IBD with one parent were considered selfed; (2) those with <90% IBD with both parents were classified as recombinants; and (3) recombinant progeny sharing ≥90% IBD with each other were defined as non-unique clones.

| Study | Cross | No. of Clones | Progeny Types |  |  | No. of Unique Recombinant |
| --- | --- | --- | --- | --- | --- | --- |
|  |  |  | NF54 Selfed | NHP4026 Selfed | Recombinant |  |
| Button-Simons et al., 2021 | NF54 x NHP4026 (before 2018) <sup>a</sup> | 146 | 1 | 0 | 145 | 88 |
| Amambua-Ngwa et al., 2023 | NF54 x NHP4026 (2019) | 158 | 2 | 0 | 156 | 113 |
| Unpublished | NF54<sup>mCherry</sup> x NHP4026<sup>GFP</sup> (2021) | 72 | 2 | 1 | 69 | 69 |

a. Multiple cloning attempts (between 2013 and 2018) of the same genetic cross conducted in 2013.

We designed experiments to better understand mating patterns of malaria parasites within mosquitoes in controlled genetic crosses where both inbred and outcrossed mating are possible. Previous work on parasite mating patterns in the mosquito have utilized PCR genotyping of polymorphic markers in individual oocysts dissected from the mosquito midgut (**Table 2**) (Ranford-Cartwright, et al. 1993; Babiker, et al. 1994; Paul, et al. 1995; Razakandrainibe, et al. 2005; Morlais, et al. 2015; Soontarawirat, et al. 2017). This is labor intensive limiting the sample size possible; studies to date have examined 35 – 623 oocysts. Furthermore, because oocysts are small and difficult to dissect without rupture, PCR methods are prone to PCR artifacts – contamination from ruptured oocysts may result in over estimation of heterozygosity, while null alleles or allelic drop out can result in underestimation of heterozygosity (Anderson, Paul, et al. 2000). Previous studies have revealed patterns ranging from random mating and good fit to Hardy-Weinberg expectations (Ranford-Cartwright, et al. 1993), to high levels of inbreeding with deficits of heterozygous oocysts and positive inbreeding coefficients (Babiker, et al. 1994; Paul, et al. 1995; Razakandrainibe, et al. 2005; Soontarawirat, et al. 2017) (**Table 2**).

**Table 2.** Summary of previous genotyping studies of oocysts (mosquito stage).

| Study | Location | No. of Oocyst | Study details | $F_{IS}$ (inbreeding in mosquitoes) | $F_{IT}$ (inbreeding in total population) | Heterozygote deficit |
| --- | --- | --- | --- | --- | --- | --- |
| Ranford-Cartwright et al., 1992 | Laboratory | 98 | Cross between 3D7 and HB3 | nd | -0.143 | No |
| Babiker et al., 1994 | Tanzania | 71 | Field caught mosquitoes | nd | 0.389 | Yes |
| Paul et al., 1995 | New Guinea | 128 | Field caught mosquitoes | nd | 0.915 (0.835-0.966) <sup>a</sup> | Yes |
| Anthony et al., 2007 | Tanzania | 35 | Field caught mosquitoes | 0.16 (0.10-0.22) | nd | Yes |
| Annan et al., 2007 | Cameroon and Kenya | 221 | Field caught mosquitoes | 0.18 (0.14-0.23) for *A. gambiae*, 0.23 (0.17-0.28) for *A. funestus* | 0.47 (0.43–0.51) for *A. gambiae*, 0.60 (0.57–0.64) for *A. funestus* | Yes |
| Razakandrainibe et al., 2005 | Kenya | 553 | Field caught mosquitoes | 0.147 (0.062-0.209) | 0.43 (0.37-0.48) | Yes |
| Morlais et al., 2015 | Laboratory | 623 | Mosquitoes infected with blood donors | 0.28 (0.234 - 0.339) | nd | nd |
| Soontarawirat et al., 2017 | Thailand | 131 | Mosquitoes infected with *P. vivax* infected blood through membrane feeder | 0.55 | 0.89 | Yes |

a. Technical issues may result in underestimation of heterozygote frequency and overestimation of $F_{IS}$ . *nd*, no data. Numbers in brackets indicate 95% confidence intervals.

In the current study, we introduced fluorescent markers into two parental parasites at the same genetic locus, so oocysts resulting from the selfing of parental parasites are labelled with mCherry (“red”), in the case of the African parasite NF54 or GFP (“green”), in the case of the Thai parasite NHP4026, and heterozygous oocysts, which carry both markers, are labeled with both fluorophores and are thus “orange”. With these marked parasites we could rapidly score genotype and size of thousands of oocysts from mosquitoes between 7- and 14-days post infection with gametocytes, allowing a detailed quantification of the dynamics of the parasite mating patterns and oocyst development. Our data reveal (i) decreasing proportions of heterozygous oocysts and thus (ii) increasing inbreeding coefficients over the course of midgut infections in two independent crosses, consistent with increased development rates of outbred oocysts and consequently, an outbreeding advantage. We also observed a significant impact of genotype on oocyst size. NF54 oocysts were 7-40% bigger than the other oocyst genotypes (NHP4026 selfed and NF54 × NHP4026 outcrossed), suggesting that oocyst size can play a role in infective stage production and transmission.

## RESULTS

### Generation of genetic crossing using fluorescently labelled parental parasites

Transgenic parasites expressing fluorescent proteins are a powerful tool in parasitology research (Hoshizaki, et al. 2022). In this study, we used two parasite strains to generate fluorescently labeled transgenic lines. NF54 is an African strain, isolated in the 1980’s, while NHP4026 was obtained in 2007 from a patient at the Shoklo Malaria Research Unit (SMRU) clinic on the Thailand–Myanmar border. We engineered NF54 to express mCherry (red) and NHP4026 to express GFP (green), generating two distinct transgenic lines: NF54^mCherry^ and NHP4026^GFP^ (**Fig. S1 and Table S1**). The fluorophore was expressed constitutively and replaced the *Pfs47* locus in both parasite parents. These edited parasite lines were then used to perform genetic crosses.

### Genetic diversity during oocyst development within the mosquito vector

Mosquito sampling was conducted across two independent crosses, with midgut dissections and oocyst collections occurring between 8 - 13 days post-infection (DPI) for *Cross 1* and 7 - 14 DPI for *Cross 2* (**Fig. 1**). Following mosquito infections, a total of 8,540 oocysts were counted and analyzed from 509 mosquitoes, including 435 that were infected (**Table S2**). This included 156 infected mosquitoes from *Cross 1* (infection rate: 0.79) and 279 from *Cross 2* (infection rate: 0.89). The mean oocyst count per mosquito was 23 in *Cross 1* (range: 1 - 137; median: 14) and 17 in *Cross 2* (range: 1 - 62; median: 13). In both crosses, oocyst burden per mosquito declined significantly over time (**Fig. 1B**, **Fig. S2**). Prior to DPI 10, the average oocyst burden per mosquito was more than 20 oocysts; however, a significant reduction was observed after DPI 10 (*Cross 1*: *p* = 4.36 × 10^−5^; *Cross 2*: *p* = 1.42 × 10^−32^; two-tailed Welch’s *t*-test) (**Table 3**), consistent with oocyst maturation resulting in sporozoite release.

**Fig. 1.**
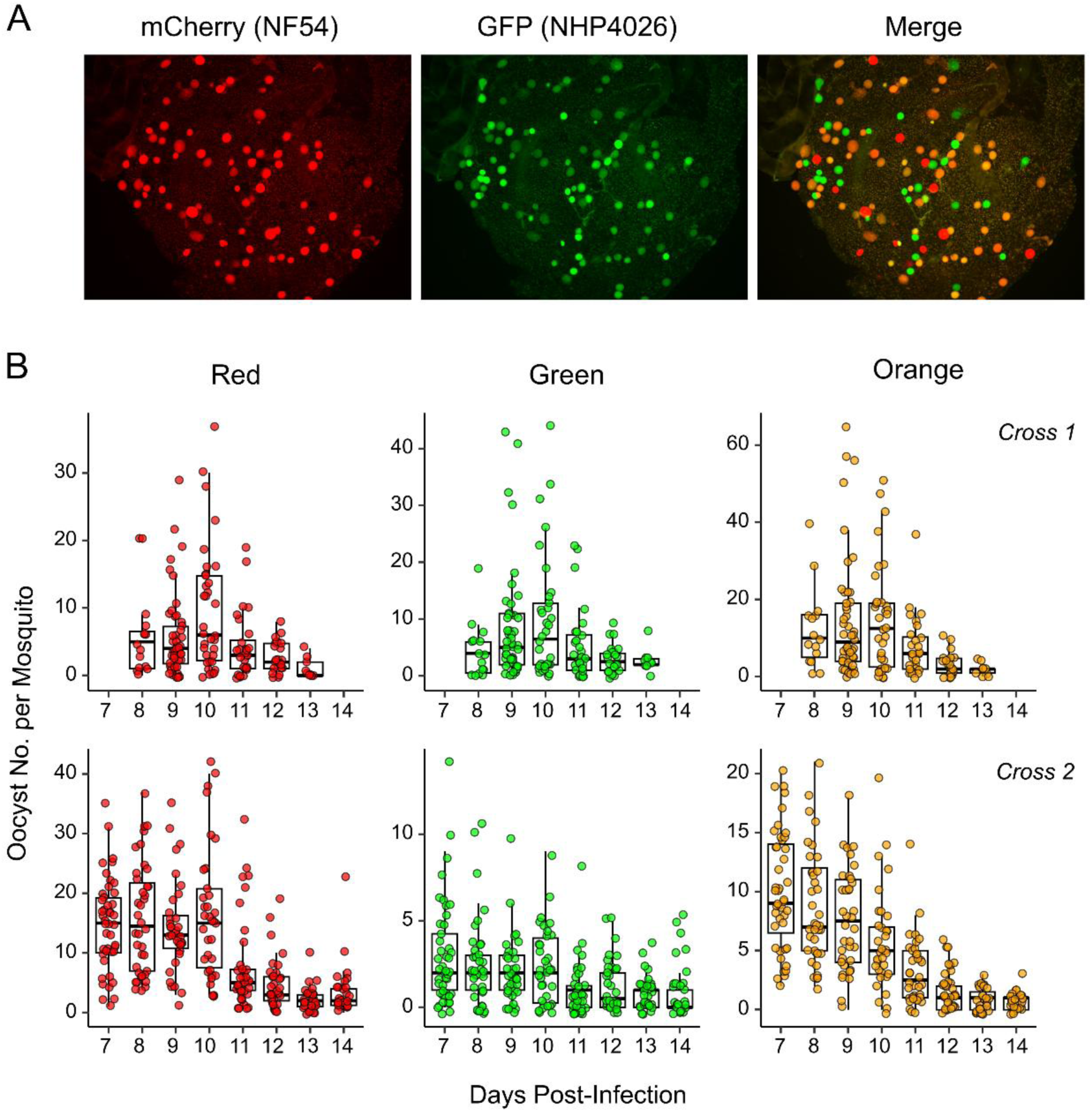
Oocysts genotype and developmental dynamics within the mosquito vector. (A) Representative fluorescence images of mosquito midguts showing oocyst genotypes. Oocysts expressing mCherry appear red, those expressing GFP appear green, and dual-labeled oocysts appear orange in the merged image. (B) Temporal dynamics of oocyst burden per mosquito across days post-infection (DPI). Panels from left to right show the number of red (mCherry-only), green (GFP-only), and orange (dual-labeled) oocysts. Top panels show data from *Cross 1*, and bottom panels show data from *Cross 2*. See Fig. S2 for plots showing total numbers of oocysts.

**Table 3.** Summary of mosquito infection data from genetic crosses between NF54^mCharry^ and NHP4026^GFP^.

| Cross | DPI | Infected/Total Mosquitoes | Total Oocyst No. | Oocyst No. per Infected Mosquito Median (Range) | HWE *p*-value | *F*-statistics (95% CI) |  |  |
| --- | --- | --- | --- | --- | --- | --- | --- | --- |
|  |  |  |  |  |  | *F<sub>IS</sub>* | *F<sub>ST</sub>* | *F<sub>IT</sub>* |
| Cross 1 | 8 | 15/17 | 342 | 21(3-68) | 0.085 | -0.110 ([-0.240]-[-0.030]) | 0.019 (0.007-0.073) | -0.089 ([-0.199]-0.013) |
|  | 9 | 48/62 | 1374 | 20.5(1-137) | 0.388 | -0.034 ([-0.102]-0.002) | 0.011 (0.014-0.040) | -0.022 ([-0.074]-0.030) |
|  | 10 | 34/36 | 1164 | 29.5(1-124) | 4.82E-06 | 0.130 (0.058-0.176) | 0.006 (0.007-0.037) | 0.135 (0.077-0.192) |
|  | 11 | 28/35 | 486 | 11.5(1-79) | 3.49E-03 | 0.115 ([-0.003]-0.182) | 0.023 (0.020-0.091) | 0.135 (0.039-0.225) |
|  | 12 | 22/31 | 195 | 9(1-19) | 8.65E-05 | 0.273 (0.078-0.351) | 0.015 (0.021-0.157) | 0.284 (0.164-0.411) |
|  | 13 | 9/17 | 52 | 4(1-17) | 0.039 | 0.322 ([-0.067]-0.548) | -0.046 ([-0.066]-0.222) | 0.291 (0.004-0.577) |
| Cross 2 | 7 | 44/45 | 1230 | 28(4-53) | 1.83E-05 | 0.111 (0.032-0.153) | 0.013 (0.016-0.051) | 0.123 (0.063-0.180) |
|  | 8 | 38/41 | 1025 | 24.5(7-57) | 2.23E-07 | 0.162 (0.082-0.213) | 0.000 (0.005-0.038) | 0.162 (0.100-0.229) |
|  | 9 | 36/37 | 862 | 22(2-53) | 1.88E-05 | 0.146 (0.054-0.199) | 0.001 (0.008-0.045) | 0.146 (0.072-0.219) |
|  | 10 | 34/35 | 845 | 23.5(4-62) | 5.74E-19 | 0.291 (0.190-0.348) | 0.023 (0.026-0.079) | 0.307 (0.231-0.382) |
|  | 11 | 36/39 | 439 | 8(1-48) | 2.03E-06 | 0.214 (0.062-0.280) | 0.018 (0.030-0.121) | 0.228 (0.123-0.337) |
|  | 12 | 34/41 | 261 | 7(1-27) | 1.03E-12 | 0.421 (0.238-0.505) | 0.039 (0.045-0.184) | 0.444 (0.329-0.560) |
|  | 13 | 31/39 | 119 | 3(1-11) | 2.15E-08 | 0.490 (0.132-0.577) | 0.053 (0.090-0.350) | 0.517 (0.335-0.675) |
|  | 14 | 26/34 | 146 | 4(1-28) | 1.67E-15 | 0.657 (0.437-0.762) | 0.011 (0.012-0.285) | 0.661 (0.517-0.802) |

#### Reduction in heterozygosity proportion among oocysts over time

We observed a significant decline in the proportion of heterozygous (outcrossed) oocysts isolated from the mosquito midgut over time, coinciding with a reduction in overall oocyst counts (**Fig. 2A**, **Table 3, Fig. S3**). In *Cross 1*, the frequency of heterozygous (orange) oocysts decreased from 0.54 on DPI 8 to 0.33 on DPI 13 (*p* = 4.84 × 10 ^−4^, linear regression). A similar trend was observed in *Cross 2*, where heterozygous oocysts dropped from a frequency of 0.36 on DPI 7 to 0.13 on DPI 14 (*p* = 3.10 × 10 ^−4^).

**Fig. 2.**
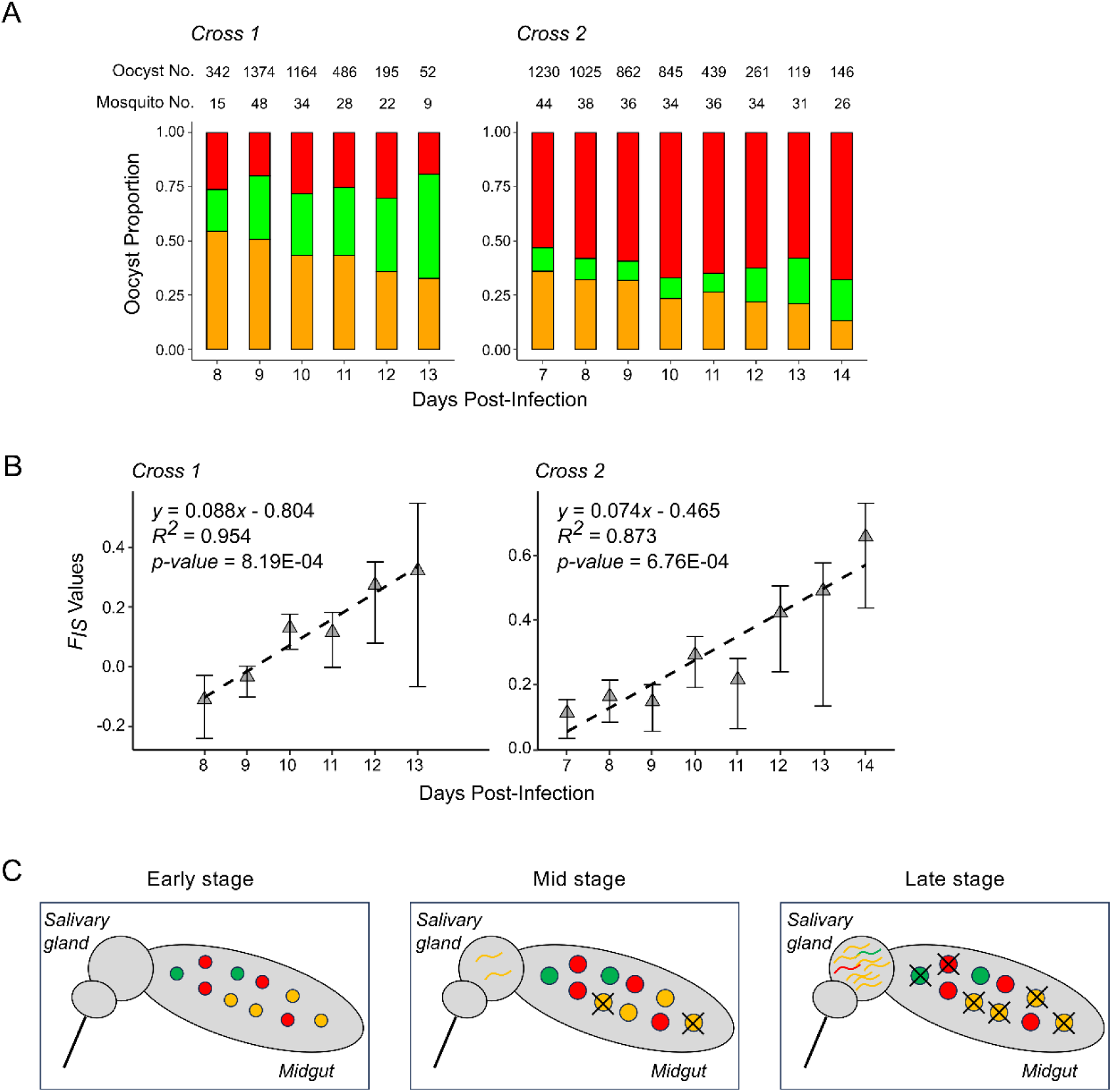
Genetic diversity during oocyst development within the mosquito vector. (A) Temporal dynamics of oocyst genotypes, shown as the proportions of red (mCherry-only, top bars), green (GFP-only, middle bars), and orange (dual-labeled, bottom bars) oocysts across days post-infection (DPI). The number of infected mosquitoes and the total oocyst count for each DPI are indicated above the corresponding panel. We observed significant reduction in proportions of outcrossed oocysts over time in both crosses (Cross 1: R^2^ = 0.964, *p* = 4.84E-4; Cross 2: R^2^ = 0.902, *p* = 3.10E-4; see also Fig. S3). (B) Regression analysis of inbreeding coefficient (*F*_IS_) over time, with 95% confidence intervals indicated by vertical lines. (C) Model illustrating how faster maturation of heterozygous oocysts may lead to an outcrossing advantage.

For both crosses, the total proportion of homozygous oocysts (red plus green) decreased over time, consistent with the increased proportion of heterozygous (orange) oocysts. However, the proportion of red or green oocysts differed between the two genetic crosses. Higher proportions of NF54^mCherry^ selfed (red) oocysts were observed in *Cross 2* but not in *Cross 1*. In *Cross 1*, the proportion of red oocysts remained relatively stable across time points (frequency from 0.26 to 0.19, *p* = 0.854), while the proportion of NHP4026^GFP^ selfed (green) oocysts increased significantly from 0.19 to 0.48 (*p* = 0.011). In *Cross 2*, NF54^mCherry^ selfed oocysts consistently dominated the population, with a non-significant increase from 0.53 to 0.68 over time (*p* = 0.093). Meanwhile, NHP4026^GFP^ selfed oocysts increased significantly from 0.11 to 0.19 (*p* = 0.019).

#### Increased inbreeding coefficient over time in mosquito midgut

To quantify the observed decline in heterozygosity proportion, we calculated hierarchical Wright’s F-statistics for each cross and time point (**Fig. 2B**, **Table 3**). The inbreeding coefficient (*F_IS_*), which reflects deviations from random mating within mosquito midguts, increased significantly over time for both crosses. In *Cross 1*, *F_IS_* rose from –0.110 on DPI 8 (95% CI: –0.240 to 0.030) to 0.322 on DPI 13 (95% CI: – 0.067 to 0.548), with a strong linear trend (*p* = 8.19 × 10 ^−4^, R² = 0.954, linear regression). Similarly, in *Cross 2*, *F_IS_* increased from 0.111 on DPI 7 (95% CI: 0.032 to 0.153) to 0.657 on DPI 14 (95% CI: 0.437 to 0.762) (*p* = 6.76 × 10 ^−4^, R² = 0.873).

In both crosses, the among-host differentiation (*F_ST_*) remained close to zero across time points, indicating little genetic structure among individual mosquitoes. Consequently, the total inbreeding coefficient (*F_IT_*) followed a similar trajectory to *F_IS_*, reinforcing the conclusion that temporal increases in homozygosity were primarily driven by within-mosquito processes rather than among-mosquito differentiation.

### Oocyst size dynamics

To investigate the factors influencing oocyst size, we performed multiple linear regression using DPI, oocyst genotype, and oocyst burden (total oocyst number per mosquito) as predictors. The analysis was conducted separately for each genetic cross (**Fig. 3, Fig. S4, Table S3**).

**Fig. 3.**
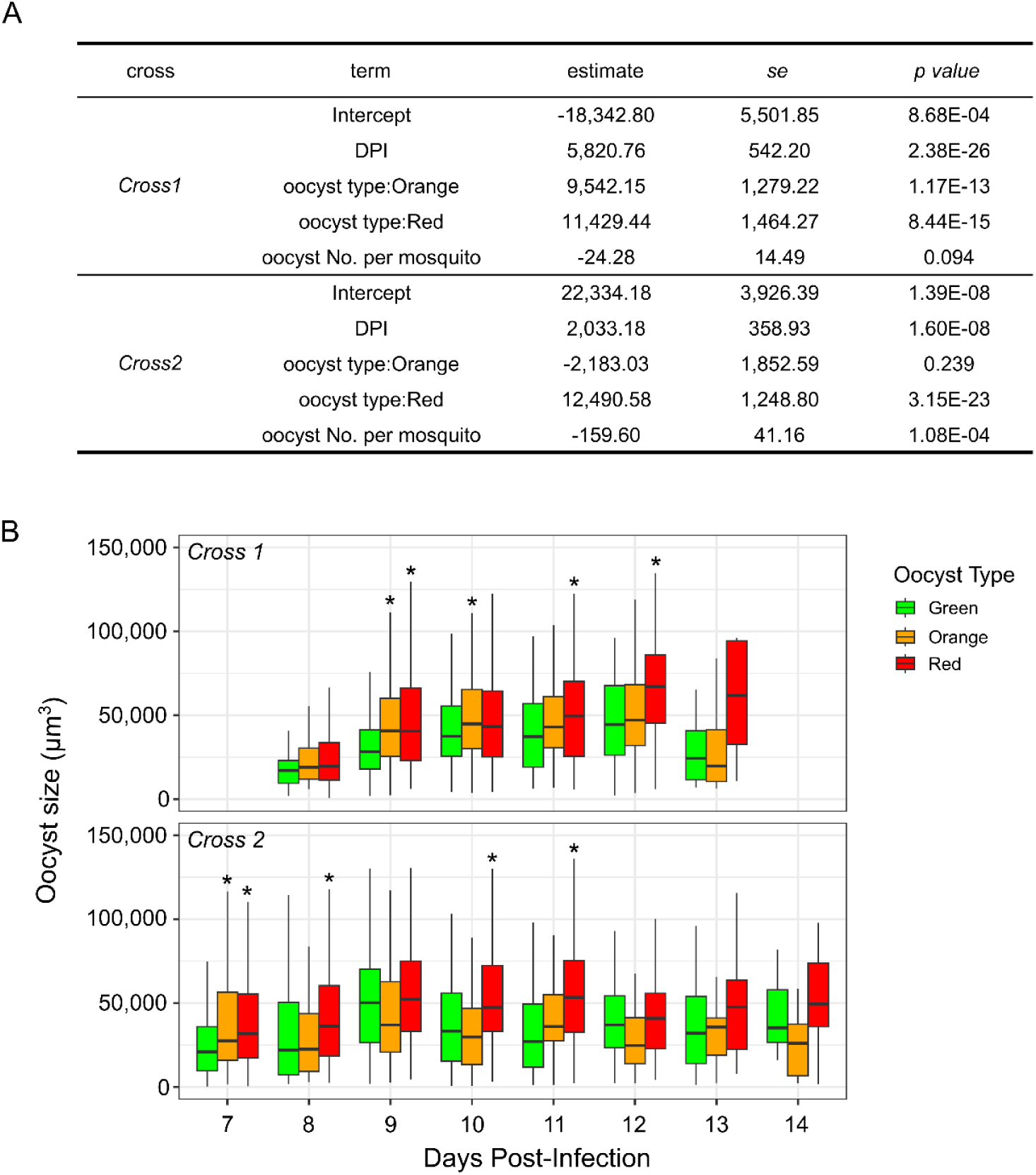
Analysis of factors influencing oocyst size. (A) Multiple regression analysis assessing the effects of oocyst genotype, days post-infection (DPI), and oocyst burden per mosquito on oocyst size across genetic crosses. (B) Box plots showing oocyst size variation by DPI and oocyst genotype across genetic crosses. For each DPI, boxes are ordered from left to right as green (NHP4026^GFP^ selfed), orange (outcrossed), and red (NF54^mCherry^ selfed). Asterisks indicate significant differences compared to the green group (*p* < 0.05).

Across both crosses, DPI was positively and significantly associated with oocyst size (volume), which is to be expected as oocysts grow larger as they mature over time. In *Cross 1*, the effect was substantial (*p* = 2.38 × 10^−26^, multiple linear regression), while in *Cross 2* the growth rate was more modest but still highly significant (*p* = 1.60 × 10^−8^). These results confirm a strong temporal component in oocyst development.

Oocyst genotypes also had a significant impact on size. NF54^mCherry^ selfed (red) oocysts were consistently and significantly larger than NHP4026^GFP^ selfed (green) in both crosses (*Cross 1*: 32.9% larger, *p* = 8.44 × 10^−15^; *Cross 2*: 40.1% larger, *p* = 3.15 × 10^−23^). Orange (heterozygous) oocysts were significantly larger than green in *Cross 1* (24.6% larger, *p* = 1.17 × 10^−13^) but showed no significant difference in *Cross 2* (*p* = 0.239). This pattern reinforces the conclusion that the NF54 (red) genotype is associated with increased oocyst size, while the effect of the heterozygous (orange) genotype may be context dependent.

Total oocyst burden showed a negative relationship with oocyst volume in both crosses. In *Cross 2*, this association was statistically significant (*p* = 1.08 × 10^−4^), suggesting a crowding effect where high parasite loads lead to reduced oocyst size, likely due to competition for space or resources. In *Cross 1*, the trend was similar but not statistically significant (*p* = 0.094).

## DISCUSSION

Fluorescent labelling of parental parasites enabled high-throughput, real-time analysis of oocyst genotype (selfed vs. outcrossed) without PCR-based artifacts in thousands of oocysts. We observed a significant temporal decline in the proportion of heterozygous oocysts within the mosquito midgut, accompanied by a corresponding increase in the inbreeding coefficient (*F_IS_*). This pattern was consistent in two independent crosses and suggests that outcrossed (heterozygous) oocysts mature and rupture earlier than homozygous (selfed) oocysts (**Fig. 2C**). Consequently, outcrossed oocysts become progressively underrepresented in the midgut at later time points due to more rapid development. Based on these dynamics, we infer that sporozoites derived from heterozygous oocysts are likely to reach the salivary glands earlier than those from homozygous oocysts, thereby gaining a temporal advantage for transmission. Faster migration of sporozoites to the salivary glands is likely to be important for transmission, given that many mosquitoes die before taking a second (infectious) blood meal. This makes mosquito longevity critical for malaria transmission models (Smith and McKenzie 2004). The observed >20% reduction in the frequency of outcrossed oocysts between days 7 and 14 post-infection corresponds to an estimated selection coefficient of ∼3 - 4% per day (**Table 3**), highlighting a substantial advantage for outcrossed oocysts in the transition to infectious sporozoites. Our data is consistent with a strong outcrossing advantage in the maturation of oocysts in the NF54^mCherry^ × NHP4026^GFP^ genetic cross.

Although our current study focused on the mosquito stages of the malaria parasite life cycle, there is evidence that the advantages of outcrossing extend to other developmental stages. In our previous work on *P. falciparum* genetic crosses between MKK2835 and NHP1337 (Li, et al. 2019), we observed strong selection against selfed progeny in pooled samples during early blood stage development, with recombinant progeny outcompeting inbred parasites carrying parental genotypes. These findings suggest that outcrossed progeny may survive and propagate more successfully in the vertebrate host. Combined with our current observation that the heterozygous oocysts in our crosses mature earlier and likely invade the salivary glands sooner, outcrossing may confer broader fitness advantages across the parasite’s life cycle. Given that liver stage development represents a critical bottleneck, future work could investigate whether similar selection pressures favor outcrossed parasites during hepatic schizogony, potentially shaping host colonization dynamics and transmission efficiency.

We suggest two possible explanations for the difference in development and maturation of inbred and outcrossed oocysts in the mosquito midgut. First, outcrossed oocysts may more effectively evade the mosquito immune response than inbred oocysts leading to faster development and rupture. Mosquito immune responses against oocysts are mediated by midgut stem cells and enteroblasts or by hemocytes (Barletta, et al. 2025) and the specific oocyst molecular signals used by the mosquito to recognize oocysts are not currently known. The mechanisms that might underlie differential survival of inbred and outbred oocysts are therefore unclear. Second, outcrossing may be advantageous because recombination between different parasite genotypes allows removal of “genetic load” (Bertorelle, et al. 2022) caused by deleterious mutations accumulated during asexual multiplication. These deleterious mutations cannot be removed during selfing because they are present in both male and female gametes. The burden of deleterious mutations may be considerable. After initial infection by mosquito inoculation, sporozoites establish infection in liver cells, where they divide rapidly. Blood stage infection in the human host is initiated by the emergence of > 90,000 clonal merozoites from infected liver cells (Vaughan, et al. 2012; Vaughan and Kappe 2017). Populations of malaria-infected red blood cells within an infected individual can reach 10^11^-10^12^. This equates to a minimum of 37-40 mitotic cell divisions if we assume geometrical cell division; in reality cell division is asynchronous (Arnot, et al. 2011) so actual numbers of cell divisions may be considerably higher. Given SNP mutation rates of 2.10-5.23 × 10^−10^ per site per asexual cycle in *P. falciparum* (Bopp, et al. 2013; Hamilton, et al. 2017; McDew-White, et al. 2019), indel and microsatellite mutations that are 3-4 orders of magnitude higher (Hamilton, et al. 2017; McDew-White, et al. 2019), and conservative estimates of 37-40 cell divisions in the human host, we estimate that on average, in mature infections, each blood stage parasite contains at least 0.45 ± 0.007 SNPs and 3,433 ± 1,591 indel/microsatellite mutations.

Single cell approaches now allow direct quantification of mutations within infections (Nkhoma, et al. 2020; Dia and Cheeseman 2021; Dia, et al. 2021). Dia *et al*. (Dia, et al. 2021) used single cell genome sequencing of natural *P. falciparum* and *P. vivax* infections. In 15 *P. falciparum* infections they identified 203 *de novo* mutations in 181 genes, while in 11 *P. vivax* infections they identified 159 *de novo* mutations in 90 genes. In both parasite species, Dia *et al*. (Dia, et al. 2021) observed strong evidence for selection, with excess non-synonymous mutations and enrichment of mutations in some gene families, including the *ApiAP2* gene family. These data are consistent with substantial genetic load resulting from accumulated mutations. Furthermore, the strong signatures of selection and non-random distribution of mutations are consistent with short term adaptation (Didelot, et al. 2016) for blood stage growth, which may have deleterious impact on parasite development in the mosquito or elsewhere in the parasite lifecycle. Outcrossing allows parasites to remove mutations that have a deleterious effect on oocyst growth and maturation, while inbreeding retains such mutations. The ookinete and oocysts are the only diploid stages of the mosquito lifecycle. Outcrossed ookinetes or oocysts are therefore more likely than inbred parasites to have fully functional gene product from at least one parent for genes critical for parasite development in the mosquito. While we suspect that removal or masking of genetic load is the principal driver of the outcrossing advantage observed in these laboratory crosses, it is also the case that in nature recombination has additional benefits for parasites. For example, recombination generates diversity in surface antigens that are key to immune evasion and allows evolution of multidrug resistance.

Malaria parasite populations range from predominantly outcrossed, to highly inbred, and this scales with levels of transmission (Anderson, Haubold, et al. 2000; Nkhoma, et al. 2013; Schaffner, et al. 2023). In countries with low levels of transmission and elevated inbreeding (e.g., South America and Southeast Asia), parasites are expected to accumulate substantial levels of genetic load and inbreeding depression, due to serial inbreeding across multiple parasite generations. Population genomic studies reveal parasites sampled 14 years apart that are identical-by-descent (IBD) in Colombia (Taylor, et al. 2020), 5 years apart in Myanmar (Li, et al. 2025) or 6 years apart in Senegal (Redmond, et al. 2018). Furthermore, in the Senegalese study (Redmond, et al. 2018) numbers of mutations increase within clonal lineages over time. Outcrossing may be help restore fitness in parasite lineages where serial sefling and inbreeding predominates. Observations that are consistent with this include: (i) multiple clone infections typically contain related parasites from recent recombination events, even in low transmission regions (Nkhoma, et al. 2012; Nkhoma, et al. 2020) and (ii) studies of genomic epidemiology in low transmission regions often contain large numbers of closely related parasites, generated from recombination (Wasakul, et al. 2023; Li, et al. 2025).

Prior studies based on PCR genotyping of oocysts in field collected mosquitoes have typically revealed mild to severe deficits of heterozygotes (**Table 2**). This pattern is consistent with our observations of heterozygote deficit in laboratory crosses and suggests that the heterozygote advantage may also drive patterns observed in the field. It is not possible to determine time since infection in field collected mosquitoes. This may influence the wide range of heterozygote deficits observed in prior studies, with older infections showing greater heterozygote deficit than young infections. Other factors may contribute to deviations from Hardy-Weinberg expectations. Hardy-Weinberg genotype ratios are expected when mating is random. However, imbalances in male/female gametocyte ratios between parental strains may skew mating outcomes away from the Hardy-Weinberg equilibrium (**Fig. S5**) (Waples 2015). Indeed, there is an extensive literature describing deviations from one-to-one gamete sex ratios in malaria parasites, and changes in sex ratio during the course of infection (Reece, et al. 2008).

Our results demonstrate a strong genetic influence on oocyst size, a key indicator of parasite developmental fitness and transmission potential (Qi, et al. 2015; Oke, et al. 2024). Contrary to expectations that outcrossed oocysts might exhibit superior growth, we observed no consistent size advantage among heterozygotes. Instead, oocysts derived from the NF54 parent were significantly larger than those from NHP4026 with heterozygous oocysts displaying intermediate sizes. Our findings are consistent with previous studies in rodent malaria parasites *Plasmodium yoelii* (Qi, et al. 2015) and *Plasmodium berghei* (Balakrishnan, et al. 2025), which also demonstrated that oocyst size is a genotype-dependent trait. In *P. yoelii*, disruption of the D-type small-subunit rRNA gene (*D-ssu*) resulted in significantly smaller oocysts and a complete loss of sporozoite development. In *P. berghei*, a chloroquine-resistance-transporter-like (CRTL) protein was shown to be essential for both oocyst growth and sporogony. These studies underscore the importance of specific genetic loci in regulating mosquito-stage parasite development. Furthermore, recent work has shown that the nutritional status of the mosquito does not fully account for variation in oocyst size, further supporting the primacy of genotype in determining this trait (Oke, et al. 2024). Together, these results reinforce the conclusion that oocyst size is a heritable, genotype-dependent phenotype shaped by parasite genomic background. The large size of NF54-derived oocysts may reflect natural variation in reproductive strategy among *P. falciparum* isolates. With over 237 genes estimated to be expressed during the oocyst stage (Howick, et al. 2019; Real, et al. 2021), future genetic mapping efforts could help identify the loci underlying this variation and provide insight into the evolutionary forces shaping transmission-related traits.

Our study has some limitations. To visualize and precisely quantify parasite genotypes, we examined a single parasite cross (NF54 × NHP4026) for which fluorescent markers were developed. Examining levels of heterozygote advantage in additional crosses to better understand the importance of this phenomenon is ultimately required. Also, while informative, fluorescent color does not capture more subtle genomic contributions (e.g., recombination patterns, epistasis). Given the unusual pattern of maternal inheritance observed in prior crosses (Li, et al. 2019; Button-Simons, et al. 2021), fluorescent markers for monitoring mitochondria and apicoplast genomes could also be informative for future work. Nevertheless, the current study clearly demonstrates that outcrossed parasites develop faster than selfed parasites and dominate early stage infections. Our results from mosquito stage infections parallel the selective advantage of outcrossed blood stage parasites relative to selfed parasites and indicate advantages to outcrossing in both the vertebrate and invertebrate host (Li, et al. 2019). These dynamics maximize the impact of limited recombination and maintain genetic diversity and adaptive potential in natural *P. falciparum* populations.

## MATERIALS AND METHODS

### Ethics approval and consent to participate

The study was performed in strict accordance with the recommendations in the Guide for the Care and Use of Laboratory Animals of the National Institutes of Health (NIH), USA. To this end, the Seattle Children’s Research Institute (SCRI) has an Assurance from the Public Health Service (PHS) through the Office of Laboratory Animal Welfare (OLAW) for work approved by its Institutional Animal Care and Use Committee (IACUC). All the work carried out in this study was specifically reviewed and approved by the SCRI IACUC.

### Generation of fluorescent-labelled parasites

We generated transgenic *P. falciparum* parasites by integrating fluorescent markers into NF54 and NHP4026 strains using a CRISPR/Cas9-based transgenesis (**Fig. S1**). Parasites were maintained as asexual blood-stage cultures following standard protocols in complete RPMI medium supplemented with either 0.5% AlbuMAX™ II (Thermo Scientific) or 10% (v/v) type O+ human serum, with daily medium changes. Fluorescent labeling was achieved through double-crossover homologous recombination, using the plasmids pLf0115 (a modified version of pLf0116 (Mogollon, et al. 2016) with EF1α-GFP replaced by EF1α-mCherry) and pLf0019 (Mogollon, et al. 2016). Transfected NF54 and NHP4026 parasites were selected with WR99210 (8 nM) and blasticidin (2.5 µg/mL) for five days, followed by cloning by limiting dilution. Successful integration was confirmed by genotyping PCRs, demonstrating disruption of the *Pfs47* gene and insertion of EF1α-mCherry in NF54 and EF1α-GFP in NHP4026 (**Fig. S1**). Oligonucleotide primers used for transfection and genotyping are listed in **Table S1**. Clones NF54^mCherry^ 2B5 and NHP4026^GFP^ 1F were used for subsequent experiments.

### Preparation of genetic crosses

We used *Anopheles stephensi* mosquitoes and human liver-chimeric FRG huHep mice, following the protocol described by Vaughan et al. (Vaughan, et al. 2015), to generate two genetic crosses (*Cross 1* and *Cross 2*) between NF54^mCherry^ and NHP4026^GFP^ parasites. Gametocytes from each parental strain were diluted to 0.5% gametocytemia in a human serum/erythrocyte mixture to prepare infectious blood meals (IBMs). The IBMs were mixed at an approximate 1:1 ratio and used to infect ∼450 mosquitoes per cross.

### Fluorescence microscopy

Midguts were dissected from *Anopheles* mosquitoes at 7–14 days post-infection in phosphate-buffered saline (PBS). Wet mounts were prepared on glass slides, and live-cell fluorescence microscopy was performed to detect and differentiate oocysts labeled with red (mCherry) and green (GFP) fluorescent markers. Images were acquired in red, green, and brightfield channels using a Keyence fluorescence microscope equipped with a 10× air objective. Oocysts were manually counted based on fluorescence: green fluorescence indicated GFP-positive (NHP4026) oocysts, red fluorescence indicated mCherry-positive (NF54) oocysts, and co-expression of both signals (appearing orange in merged images) was used to identify heterozygous oocysts. Counts were recorded per midgut and classified by color to quantify the proportions of homozygous and heterozygous oocysts.

### Oocyst size quantification

We quantified oocyst size using QuPath v0.2.3 (Bankhead, et al. 2017). Fluorescence microscopy images of dissected mosquito midguts were imported into the software and individual oocysts were initially annotated manually using the polygon tool to train the program for automated analysis. Subsequent automated batch analyses were then performed to classify oocysts by fluorescent label (red, green, or merged) and to calculate their area (in µm²) based on the calibrated image scale. The measured areas were then converted to volume estimates to approximate oocyst size (volume, in µm^3^), assuming that oocysts are spherical. All measurements were exported for downstream statistical analysis. Consistent imaging settings and scale calibrations were maintained across all samples to ensure accuracy and comparability.

### Statistical analysis and measurement of deviations from Hardy-Weinberg Equilibrium

To investigate the population genetic structure of oocysts, we estimated Wright’s *F*-statistics, which quantify deviations from expected genetic diversity under random mating. Oocyst genotypes were analyzed across three hierarchical levels: **I** – individual oocysts, **S** – subpopulations of oocysts within individual mosquitoes, and **T** – the total population of oocysts across all mosquitoes.

The *F*-statistics were calculated as follows:

- 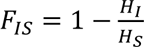

*F_IS_* ranges from −1 to 1 and measures inbreeding coefficient of individual oocysts relative to individual mosquitoes. A value of 0 indicates random mating, negative values indicate an excess of heterozygotes consistent with outcrossing, and positive values indicate a deficit of heterozygotes, suggesting selfing or inbreeding.

- 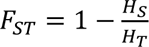

*F_ST_* ranges from 0 to 1 and measures genetic differentiation among subpopulations (individual mosquitoes). A value of 0 indicates no differentiation, while higher values reflect increasing population genetic divergence. In rare cases, *F_ST_* values may fall below zero due to statistical noise and are generally interpreted as indicating no genetic differentiation between populations.

- 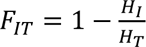

*F_IT_* ranges from –1 to 1 and quantifies the inbreeding coefficient of individuals relative to the total population. A value of 0 indicates random mating across the entire population. Negative *F_IT_* values reflect an excess of heterozygotes, suggesting high levels of outcrossing, while positive values indicate reduced heterozygosity, consistent with inbreeding or population structure.

Where:

- *H_I_* = observed heterozygosity at the individual oocyst level
- *H_S_* = expected heterozygosity within mosquitoes (subpopulations)
- *H_T_* = expected heterozygosity in the total oocyst population

These statistics were computed using the R packages *HardyWeinberg* (Graffelman 2015) and *hierfstat* (Goudet 2005). The 95% confidence intervals (CIs) were calculated using a bootstrap approach with 1,000 resampling replicates. Significance testing of the fixation indices was performed using non-parametric permutation tests to assess whether observed values significantly deviated from those expected under random mating.

## DATA AVAILABILITY

All data supporting the conclusions of this study are included in the main text and/or Supplementary Materials. Genotype data for cloned progeny are available in the following GitHub repository: https://github.com/emilyli0325/P01-cloned.progeny. Additional data related to this paper may be requested from the authors.

## AUTHOR CONTRIBUTIONS

XL, TJCA, AMV, and SK conceived and designed the study. BAA, NR, MTH, ASL, HP, AMV and SK generated the fluorescent-labeled *Plasmodium falciparum* parental lines. BAA, NR and MTH and SK carried out the genetic crosses, dissected mosquito midguts, and quantified oocysts. KSJ performed oocyst size measurements. XL conducted statistical analyses and data visualization, with input from TJCA. XL, TJCA, AMV, and SK drafted the initial manuscript. All authors contributed to critical revision of the manuscript, had full access to all data, and approved the final version for submission.

## ACKNOWLEDGMENTS

This work was supported by National Institutes of Health (NIH) program project grant P01 AI127338 (to MTF and AMV) and by NIH grant R37 AI048071 (to TJCA). Work at Texas Biomedical Research Institute was conducted in facilities constructed with support from Research Facilities Improvement Program grant C06 RR013556 from the National Center for Research Resources. The authors thank Drs. Shinya Miyazaki and Chris Janse from Leiden university for providing plasmids pLf0115, pLf0116 and pLf0019 for the generation of fluorescent parasites.

